# Combining 3D-MOT with motor and perceptual decision-making tasks: conception of a life-sized virtual perceptual-cognitive training paradigm

**DOI:** 10.1101/511337

**Authors:** Thomas Romeas, Romain Chaumillon, David Labbé, Jocelyn Faubert

## Abstract

The present study introduces a virtual life-sized perceptual-cognitive paradigm combining three dimensional multiple object tracking (3D-MOT) with motor (Experiment 1) or perceptual (Experiment 2) decision-making tasks. The objectives were to assess the impact of training on task performance and to determine the best training conditions for improvement and learning.

Seventy-one participants were randomly trained under one of four training conditions (isolated 3D-MOT task, 3D-MOT simultaneously combined with a decision-making task, consolidated 3D-MOT and decision-making task, isolated decision-making task). Task performance was evaluated using speed thresholds, decision accuracy (%) and reaction time (s).

Findings showed that the dual-task paradigm allowed satisfactory degrees of performance on both tasks despite an important dual-task cost. Interestingly, the results seemed to favor consolidated over simultaneous training for dual-task performance when 3D-MOT was combined with a motor task. The amount of attentional shared resources in regards to the nature of the additional task was discussed.

## Introduction

In the past decades, many off-field perceptual-cognitive training tools were developed to train anticipation and decision-making abilities in athletes including the 3 Dimensional-Multiple Object Tracking (3D-MOT) paradigm (Broadbent, Causer et al. 2015, Romeas, Guldner et al. 2016). However, while searching for higher performance benefits (*i.e.* far transfer), sports scientists have raised concerns in regards to the capability of these perceptual-cognitive programs. Indeed, some authors suggested that these trainings should be improved in considering broader, contextual, non-kinematic sources of information (Canal-Bruland and Mann 2015) as well as perception-action coupling (Broadbent, Causer et al. 2015).

The 3D-MOT training methodology has been developed to train the processing of visual dynamic scenes reflecting some of the fundamental and attentional demands required during sports (*i.e.* keeping track of teammates, opponents and ball). Correctly perceiving and integrating relevant information from a dynamic scene is key for an athlete when executing the most appropriate decision and action in sport. The 3D-MOT exercise requires the user to process complex motion using dynamic, sustained and distributed attention as well as working memory (Faubert and Sidebottom 2012, Parsons, Magill et al. 2016). The paradigm has been shown to be very sensitive to sport expertise and has demonstrated evidence of transfer across different populations and domains including sports (Legault and Faubert 2012, Faubert 2013, Parsons, Magill et al. 2016, Romeas, Guldner et al. 2016, Vartanian, Coady et al. 2016). The main advantages of this non-contextual technique are that it is simple to use, malleable in terms of development, allows major gains with minimal training time and can be generalized to a variety of sports or environments (Faubert 2013, Mangine, Hoffman et al. 2014, Harenberg, McCaffrey et al. 2016, Romeas, Guldner et al. 2016, Hoke, Reuter et al. 2017, Michaels, Chaumillon et al. 2017). However, this training technique lacks specificity and does not include the combination of perception and action that would be typically present in the sport environment.

Consequently, the main challenge of this study was to investigate new training conditions that would present higher fidelity with the sport requirements (*i.e.* broader contextual information) to potentially increase the transfer of learning and therefore athletic performance. Given that many sports skills involve performing multiple tasks simultaneously, performance on these skills may be enhanced if training conditions also incorporate multiple tasks (Gabbett, Wake et al. 2011). A simulated dual-task paradigm would also be closer to the attentional demand required in the sports environment (Memmert 2009). Moreover, dual-task training is often used by coaches to increase the cognitive demand of training sessions. That is why the paradigm employed in this study combines the processing of specific information with a decision-making action required while simultaneously performing the 3D-MOT task. In order to improve the ecological validity of this dual-task training, the life-sized scenario was displayed using virtual reality. As a result, the perceptual-cognitive challenge raised by this virtual dual-task paradigm was thought to be very similar to the visual and dynamic sources of information found within the sports environment.

The present study served to develop a new 3D-MOT methodology of combined perceptual-cognitive training seeking higher ecological value to respond to the field demand. The first objective of this study was to evaluate the impact of the combined training methodology on task performance. The second objective was to determine the best training condition for improvement and learning. An additional objective was to observe the impact of the nature of the decision-making task on the combined training outcome. Indeed, there is more evidence that the amount of sensory modalities shared between the combined tasks can impact attentional resources available for task performance (see Wahn and Konig 2017 for a recent review). In this regard, two experiments were conducted. The first experiment consisted of a perceptual 3D-MOT task combined with a motor decision-making exercise. The second experiment focused on a perceptual 3D-MOT task combined with a perceptual decision-making task.

## Experiment 1

A critical requirement for an athlete when competing in his sporting environment is to be able to perform multiple tasks simultaneously. For exemple, keeping track of relevant teammates, opponents or a ball while executing specific motor skills. In this regard, the first experiment consisted of a life-sized combination between 3D-MOT and a sport-specific motor task, *i.e.* birdie interception (badminton). There are a few attempts of combining MOT with motor tasks (*e.g.* locomotion) in the literature. For instance, Pothier and colleagues (2014) tested MOT while simultaneously walking with young and older adults (Pothier, Benguigui et al. 2015). They found a decrease in performance on MOT with increasing complexity of the MOT task. Importantly, an age-related decrease in MOT and gait performance was found with older adults’ performance being impaired under high attentional load conditions. This result was congruent with a previous study by Thomas and Seiffert on MOT and walking (Thomas and Seiffert 2011). This evidence raised the importance of highly challenging MOT conditions in the context of performance. One study involving professional athletes showed that training on 3D-MOT while simultaneously standing up compared to sitting-down could significantly impact 3D-MOT performance all along the training (Faubert and Sidebottom 2012). This result was corroborated by the findings of a study including Olympian-level water polo, taewkondo and tennis athletes in which participants were first trained on 3D-MOT sitting-down (consolidation), then later trained on 3D-MOT while standing-up and combined with a sport-specific position where balance was required (Quevedo, Blázquez et al. 2015). The 3D-MOT performance was systematically impacted by the addition of a secondary motor task but athletes seemed to recover their performance on 3D-MOT after a few training sessions. The study also hinted at evidence of transfer following training assessed by self-reports in athletes and coaches. However, whether the combination of 3D-MOT with an additional motor task was simultaneous (Faubert and Sidebottom 2012) or consolidated (Quevedo, Blázquez et al. 2015), it is still unclear which training methodology is more beneficial for task performance enhancement mainly because the additional task performance was not assessed in these previous studies.

In the present experiment, while we anticipate an important dual-task cost generated by the addition of a motor task to the 3D-MOT exercise, we still expect reasonable degree of performance on both exercises in addition to significant improvement throughout training sessions. However, we expect a different degree of improvement across training regimens and seek to determine the best training conditions between single, simultaneously or consolidated training for dual-task improvement. To test these hypotheses, four groups were used including different training regimens: isolated 3D-MOT task, dual-task 3D-MOT (combined training), isolated 3D-MOT task then dual-task 3D-MOT (consolidation training) and isolated decision-making task. All groups were evaluated on each task (single-task 3D-MOT, dual-task 3D-MOT, single decision-making task) three times during the training at sessions 1, 6 and 12 in order to quantify the benefits of each training regimen across time.

## Material and methods

### Participants

Twenty-nine university badminton athletes (6 women; mean = 22.98 ± 2.77 (SD) years old; 10.37 ± 4.49 years of badminton practice; 8.27 ± 3.92 hours of badminton workout per week) were recruited for the study. All participants were healthy and reported normal or corrected-to-normal vision (*i.e.* visual acuity score of 6/6 or better with both eyes in Snellen chart and stereoscopic acuity of 50 seconds of arc or better in Frisby test). Screening surveys indicated that participants were free of visual, neurological, musculoskeletal, cardiovascular and vestibular impairments. The experimental protocol and related ethics issues were evaluated and approved by the ethics committee of the Université de Montréal and were carried out in accordance with the tenets of the Declaration of Helsinki (last modified, 2004). All subjects were given verbal and written information on the study and gave their verbal and written informed consent to participate.

### Apparatus

The study was conducted using a fully immersive virtual environment (EON Icube^TM^) to give a life-sized representation of the action. The EON Icube^TM^ is a 7 × 10 × 10 feet room that includes three rigid back projection surface walls (one frontal and two laterals) and a reflective floor. Four high-resolution projectors were synchronized and the image was updated in real-time to maintain the true viewing perspective of the observer. The EON Icube™ was under the computer control of an Intel Xeon E5530 (NVIDIA Quadro FX 5800 graphic card) along with four Hewlett Packard Z800 workstations generating a stereoscopic environment. All the tasks were perfomed in the 3D environment and the stereoscopy was generated with CrystalEyes^®^ 4 s (RealD) active shutter glasses synchronized at 120 Hz. Participants were positioned at a distance of 1.2 meters from the frontal surface wall.

### Experimental setup

The three-dimensional Multiple Object Tracking (3D-MOT) task was working under the NeuroTracker™ system licenced by CogniSens Inc. (Montreal, Canada). The CORE mode of the NeuroTracker™ system was used. During the exercise, four of eight projected spheres had to be tracked within a 3D virtual volumetric cube space with virtual light grey walls, subtending a visual angle of 46 degrees (Figure 1). The spheres followed a linear trajectory in the 3D virtual space. Deviation occurred only when the balls collided against each other or the walls. Each CORE, based on a staircase procedure, lasted approximately 8 minutes. The staircase procedure consists of increasing the speed if the subject correctly identified all the indexed targets or decreasing the speed if at least one target was missed. After each correct response, the dependent variable (speed of ball displacement) was increased by 0.05 log and decreased by the same proportion after each incorrect response, resulting in a threshold criterion of 50%. Speed thresholds were then evaluated using 1-up 1-down staircase procedure (Levitt 1971) with twenty trials. CORE scores of participants corresponded to the speed threshold computed through the mean of the speeds at the last four inversions. Each participant performed three CORE in a session and completed the task while seated. Participant’s performance corresponded to the mean of the three CORE speed thresholds.

**Figure 1.**
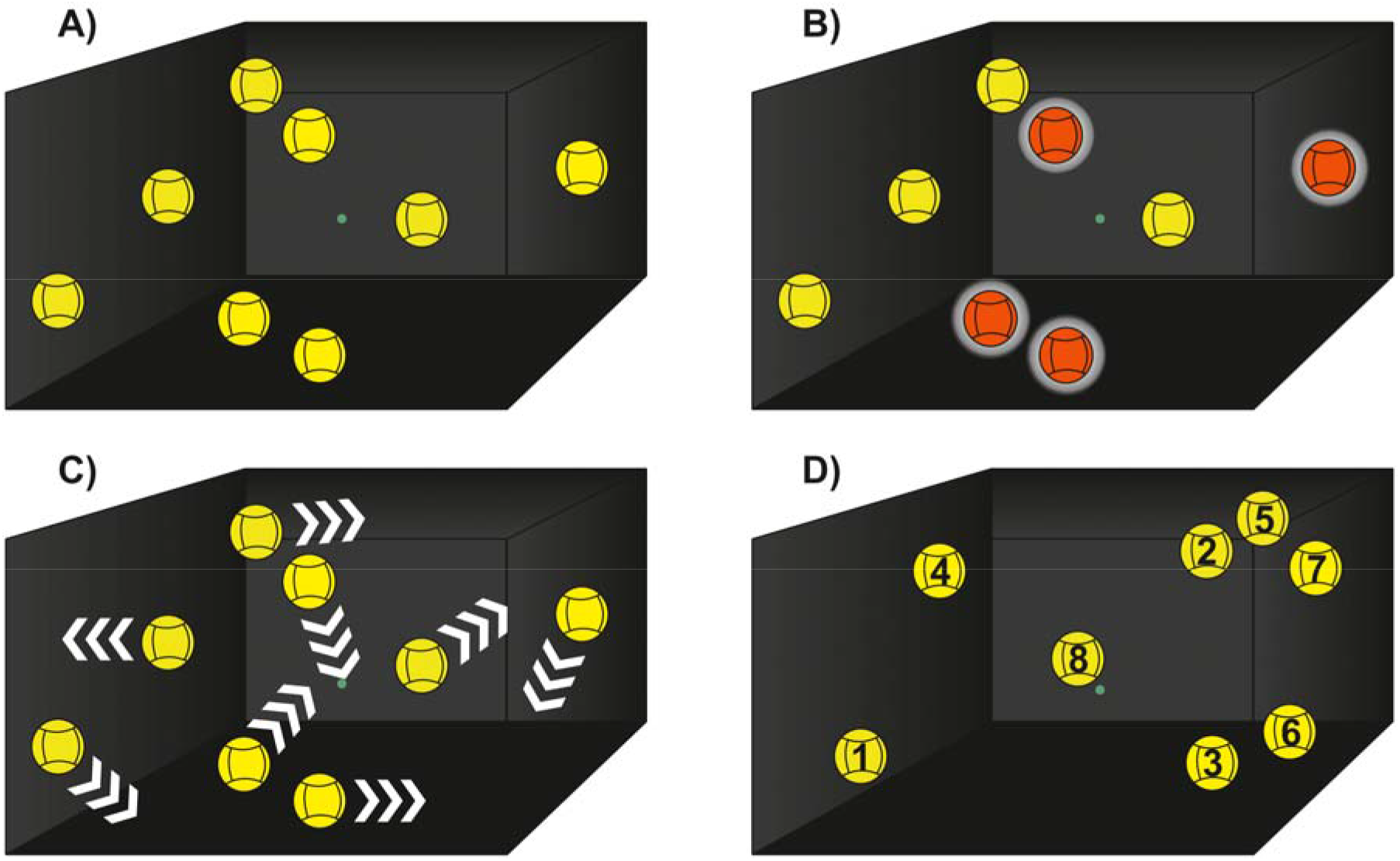
Illustration of one trial during the 3D-MOT task. A) Presentation of randomly positioned spheres in a virtual volumetric space. B) The 4 spheres to be tracked during the trial are quickly highlighted in red. C) Removal of identification and movement of all spheres with dynamic interactions. D) Observer’s response by identifying the spheres. Then a feedback is given to the observer. If the observer correctly identifies all spheres, the task is repeated at a faster speed. If, on the other hand, the observer makes a mistake, the task is repeated at a slower speed.

Our participants were badminton athletes. Therefore, the motor-response decision-making task was designed to echo their expertise and thus consisted of a birdie interception task (Figure 2). This task consisted of intercepting the trajectory of an incoming virtual birdie (virtual bird feathers length = 0.1 m; bird feathers radius = 0.07 m) with a real badminton racket, before it touched the ground. Two trajectories were implemented, and the virtual birdie could finish its race to the left or to the right of the participant (*i.e.* forced choice paradigm). These two trajectories were modeled on a real drop shot trajectory that was recorded with a 12-camera 3D motion capture system (Optitrack, NaturalPoint, Corvallis, OR). In each case, the birdie trajectory was initiated in the upper central part of the screen (*x*= 0; *y*= 2 m and *z*= 6 m). The duration of the presentation lasted 1 s and contained 60 frames. The inter-stimulus interval was 500 ms. The position of the racket was monitored with the Microsoft Kinect (V2) technology: while participants held a real racket, the racket string spatial coordinates were modeled relative to the wrist joint position which was used as a reference point for the kinect. To minimize the error rate due to mental misconception of virtual space, the string racket width and radius were configured at 0.7 and 0.75 meters, respectively. To improve the accuracy of interception point computation, the length of the arm, as well as the lateralization of the subject, were taken into account. As it is often the case in real-life badminton practice, the birdie trajectories were computed to force the subject to make one step forward to intercept the birdie before it touched the ground. Participants were instructed that they simply had to intercept the birdie without the need for a strike action. Each trajectory (*i.e.* leftward or rightward) was randomly presented thirty times in each of the three experimental blocks (a total of 180 trials). Participants were standing up during the task and their performances were assessed through their success rate (in %) and their response speed (difference in seconds between the appearance and the interception of the birdie; *i.e.* RT).

**Figure 2.**
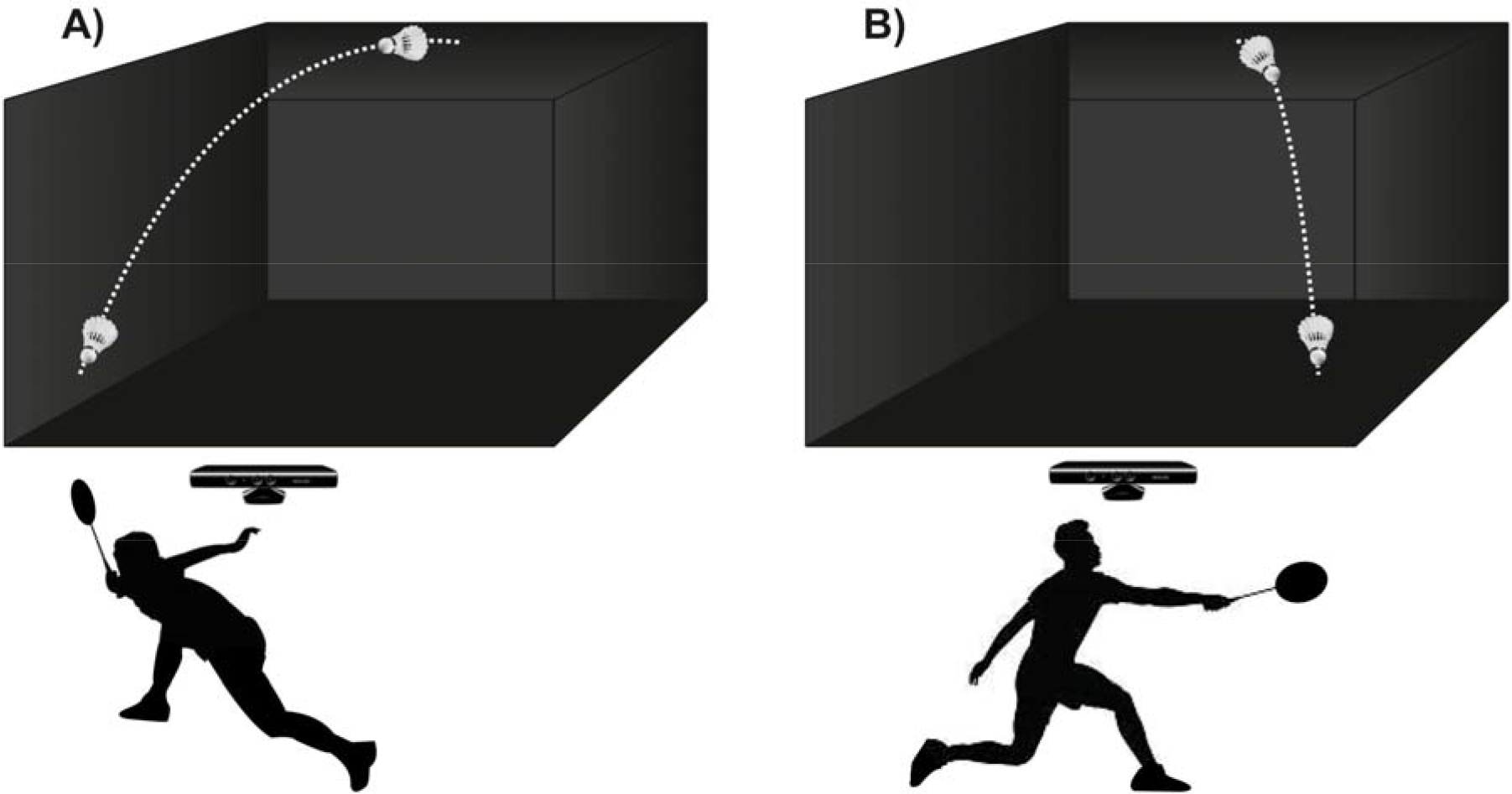
Illustration of the Birdie Interception task. The task consists of intercepting the trajectory of an incoming virtual birdie with a real badminton racket, before it touched down. Two trajectories were possible: A) leftward or B) rightward from the subjects own vertical reference. White dashed lines are used here as a visual aid and were not presented during the experiment.

### Training regimen

One objective of the present experiment was to determine the best training conditions for dual-task processing, improvement and learning. To address this question, participants were randomly trained under one out of four training regimens. *i)* Eight participants were engaged in the isolated 3D-MOT regimen (iMOT) which consisted of 3D-MOT training in its original form (Faubert and Sidebottom 2012, Faubert 2013, Romeas, Guldner et al. 2016) and served as a reference group for the standard 3D-MOT task. To test whether an additional task could impact the training, two groups were created. *ii)* the first combined training group engaged eight participants in a simultaneous dual-task training called combined (Combi) during which a combined motor task was coupled with a 3D-MOT task. During each 8 s 3D-MOT trial, participants had to simultaneously perform 3 birdie interceptions. *iii)* The second combined training group involved eight participants in a training regimen that we called consolidation (Consol). This training was designed with two different phases. The participants performed iMOT during the first four training sessions (*i.e.* phase 1, consolidation of skills on the primary task) while they performed the Combi training in the following 5 sessions (*i.e.* phase 2). *iv)* Finally, five participants performed the birdie interception task training (iDM) and served as a control group for the secondary task.

### Protocol

Regardless of their regimen, participants were trained during 9 sessions lasting 30 min (phase 1 from T02 to T05 and phase 2 from T07 to T11) and performed 3 evaluation sessions (at T01, T06 and T12) lasting one hour and a half (Figure 3). Besides the nature of the task, the experimental setup, display, duration as well as sessions frequency (twice a week for six consecutive weeks) were identical between the four training regimens. At T01, a short demonstration was given to the participants before each one of the three tasks. In the single-task 3D-MOT condition (ST_MOT_), participants were asked to reach the highest possible score. In the single-task decision-making condition (ST_DM_), instructions were to be as accurate and as quick as possible without prioritizing speed over accuracy and conversely. In the dual-task condition (DT), instructions were to equally pay attention to both the 3D-MOT and the decision-making exercise. During the evaluation sessions, all participants performed the ST_MOT_, ST_DM_ and DT conditions. Consequently, 6 performance measures were recorded at T01 (pretraining evaluation), T06 (mid-term evaluation) and T12 (post-training evaluation): speed threshold in ST_MOT_, decision accuracy and response speed in ST_DM_ as well as these same three measures in DT condition. During the training sessions, participants were engaged in their specific training.

**Figure 3.**
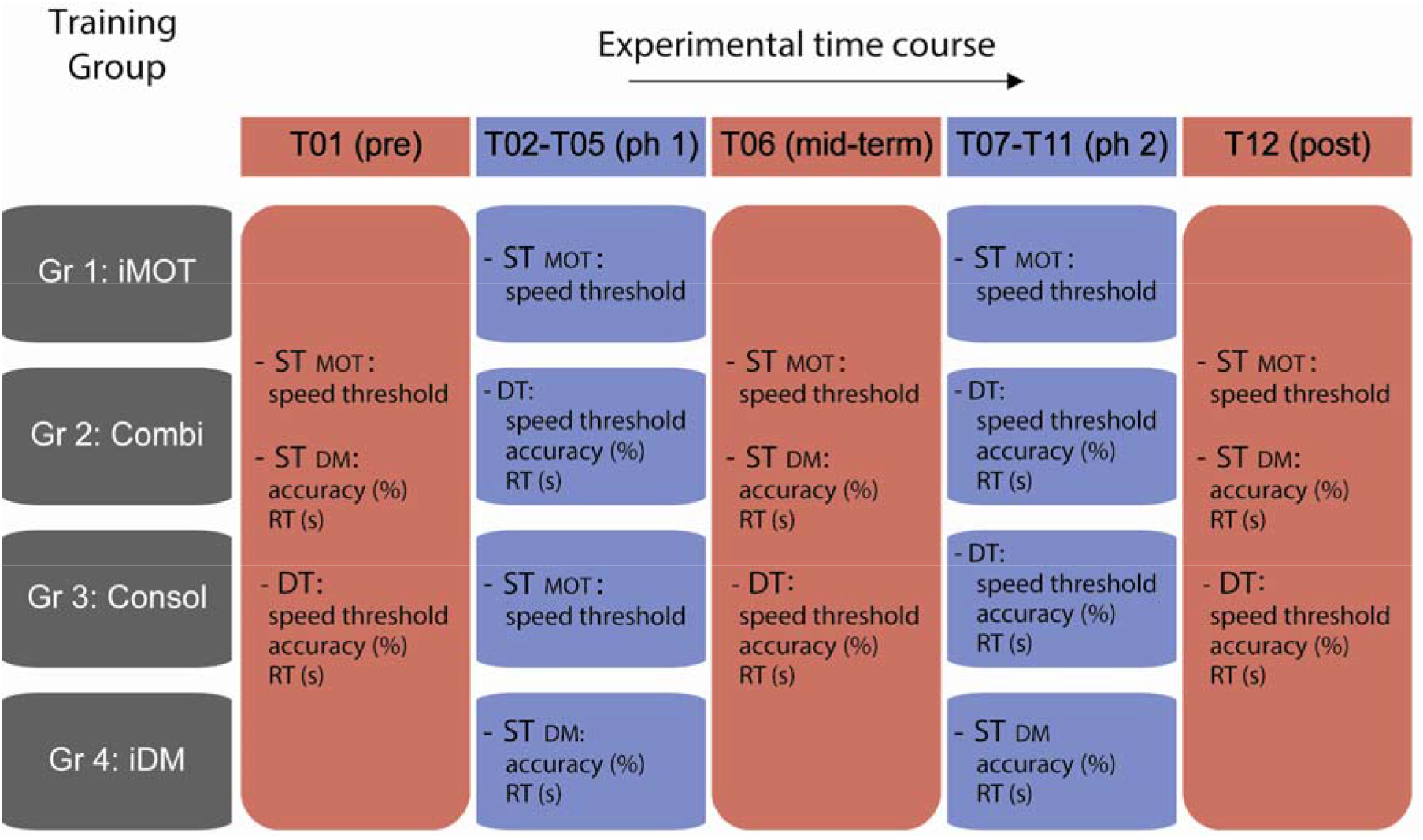
Summary of the tasks and measures performed across the 12 sessions in each of the four training regimen. Ph corresponds to phase, iMOT corresponds to isolated Multiple Object Tracking, Combi to combined, Consol to consolidation, iDM to isolated Decision-Making task, ST_MOT_ to single-task MOT, ST_DM_ to single-task decision-making and DT to dual-task.

### Statistical analyses

The objective was to test whether the combination between 3D-MOT and a specific motor task (*i.e.* dual-task training) was disruptive or beneficial for the training effectiveness and to assess the effectiveness of different training regimens across time. For each statistical analysis, eta-squared values (□^2^) were reported to provide information about the magnitude of effects. For clarity reasons, only relevant results were reported in the manuscript but a full description of the analyses’ outcome can be found in **Supplementary Material**.

Six dependent variables were used including speed threshold in STMOT and DT, decision accuracy and response speed in STDM and DT. We ensured that the variables were normally distributed by using Shapiro-Wilk tests (*p* > 0.05) and by looking at asymmetry and skewness. When the normality of distribution was ensured, analyses of variance (ANOVA) were performed and Levene’s tests were used to assess the homogeneity of variance (*p* > 0.05). Greenhouse-Geisser corrections were applied when sphericity was violated (*p* < 0.05). Multiple comparisons using Bonferroni corrections were performed to explore differences within each factor. Non-parametric tests were used when variables were not normally distributed (see details below). Note that the accuracy and response speed data from two participants (iMOT and Consol groups) were missing in session T01 due to technical errors.

To test dual-task cost and training effectiveness on it, speed thresholds, accuracy and RT from ST_MOT_, ST_DM_ and DT were submitted to a 4 (Between; *Groups*: iMOT, Combi, Consol, iDM) by 3 (Within; *Sessions*: T01, T06, T12) by 2 (Within; *Task condition*: isolated, combined) ANOVA with repeated measures on the last two factors.

To analyze the training effectiveness, each of the 6 dependent variables was submitted to a 4 (Between; *Groups:* iMOT, Combi, Consol, iDM) by 3 (Within; *Sessions:* T01, T06, T12) ANOVA with repeated measures on the last factor. To test training effectiveness across sessions, Student’s paired t-tests were used to evaluate the impact of training across *Sessions* on the six dependent variables within each training group. In addition, logarithmic regression functions were fitted on speed thresholds to observe learning rates, in particular during the DT condition. Therefore, the slope was used to compare learning performance on 3D-MOT during the first 7 sessions of DT exposure between group Combi (*i.e.* S01-S07) and group Consol (*i.e.* S06-S12). A non-parametric Mann-Whitney U test was employed. To control for a potential speed-accuracy tradeoff on the motor decision-making task, Pearson’s correlations were used to explore the relationship between RT and accuracy in STDM and DT conditions at sessions T01, T06 and T12. The test revealed no speed-accuracy tradeoff.

Finally, to test training effectiveness between groups, one-way ANOVAs with the between-subject factor *Groups* were performed separately on each of the six dependent variables in each condition and for each of the three evaluation sessions (*i.e.* T01, T06 and T12).

## Results

### Dual-task cost

There was a significant main effect of *Task condition* on speed thresholds (F[1,25] = 244.720, p < 0.001, □^2^ = 0.907) which demonstrates a change in performance between the single and combined task (Figure 4). This dual-task cost decreased with time as the analysis revealed a significant interaction between *Sessions* and *Task condition* (F[2, 50] = 6.560, p = 0.003, □^2^ = 0.208). Multiple comparisons revealed that dual-task cost was reduced between sessions T01 > T06 > T12 (all p < 0.017). There was a main effect of *Groups* (F[1, 25] = 3.116, p = 0.044, □^2^ = 0.272) and multiple comparisons showed a superior overall dual task cost in iDM compared to Consol group (p = 0.044). Regarding the motor task, there was a significant main effect of *Task condition* on RT (F[1, 23] = 25.044, p < 0.001, □^2^ = 0.521) but not on accuracy (F[1, 23] = 0.098, p = 0.758, □^2^ = 0.004). This effect demonstrates a change in RT performance with the single motor task being executed faster than when combined with 3D-MOT. There was no interaction between *Sessions* and *Task condition* (F[2, 46] = 0.292, p = 0.748, □^2^ = 0.013).

**Figure 4.**
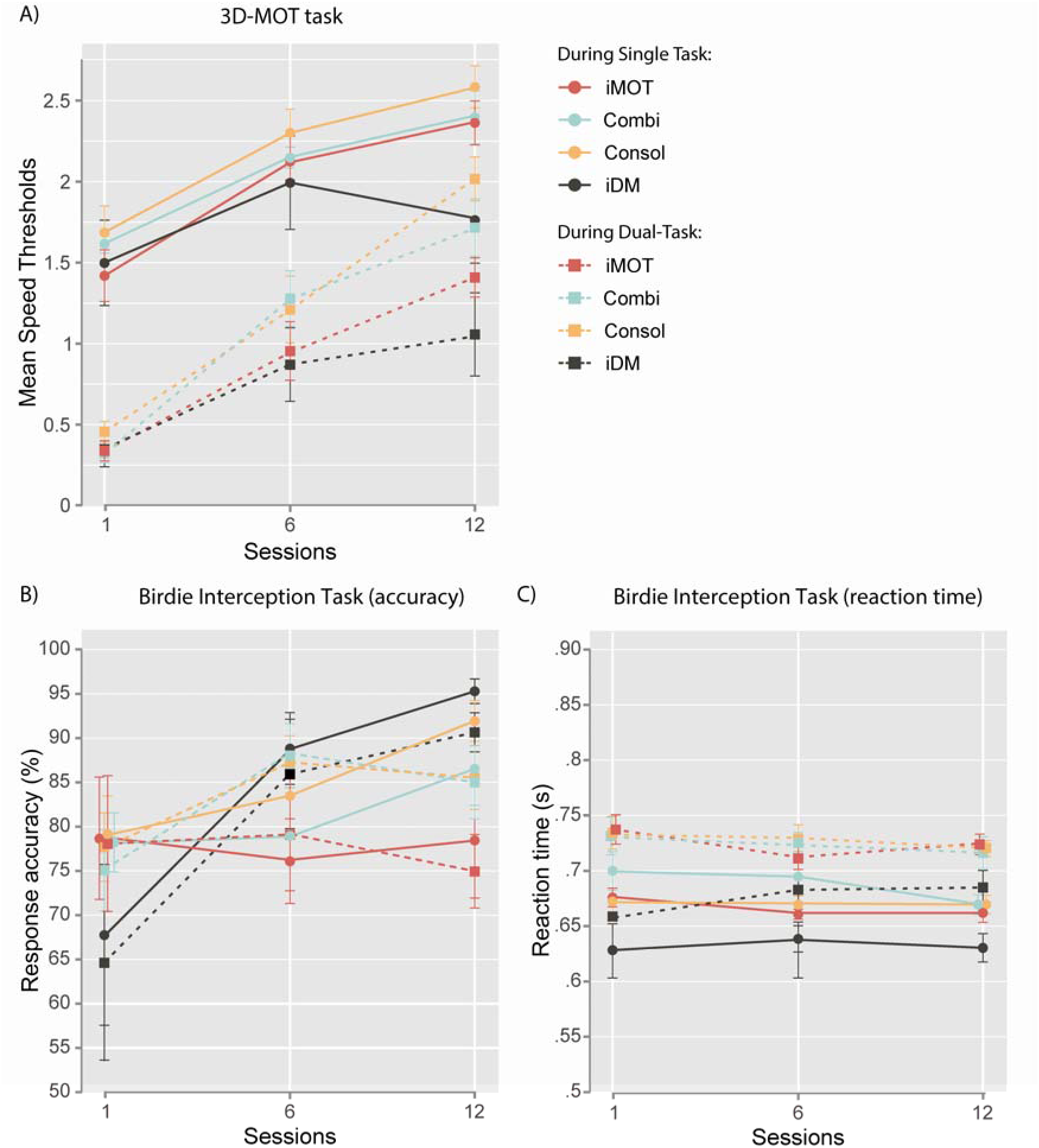
Training effectiveness across groups and sessions on the A) 3D-MOT and B), C) Decision-Making tasks (accuracy and reaction times, respectively). Circles and solid lines represent the performances obtained in single condition whereas squares and dashed lines correspond to performances recorded during the dual-task condition. Error bars represent SEM.

### Training effectiveness across Sessions and between Groups

There was a main effect of *Sessions* on speed thresholds in ST_MOT_ (F[2, 50] = 82.231, p < 0.001, □^2^ = 0.767) and DT (F[2, 50] = 86.721, p < 0.001, □^2^ = 0.776) and on accuracy in ST_DM_ (F[2, 48] = 6.298, p = 0.009, □^2^ = 0.208) and DT (F[2, 48] = 8.110, p = 0.003, □^2^ = 0.253). These effects indicate a general improvement in 3D-MOT performance and birdie interception accuracy in single and dual-task across time. There was no main effect of *Sessions* on RT in ST_DM_ (F[6, 48] = 0.515, p = 0.524, □^2^ = 0.021) or DT (F[6, 46] = 0.190, p = 0.759, □^2^ = 0.008). A significant difference between *Groups* was demonstrated in speed thresholds during DT (F[3, 25] = 4.036, p = 0.018, □^2^ = 0.326) which indicated different 3D-MOT performance between training regimens while performing the combined task. The ANOVA revealed a significant interaction between *Groups* and *Sessions* factors on speed thresholds in ST_MOT_ (F[6, 50] = 3.151, p = 0.011, □^2^ = 0.274) and DT (F[6, 50] = 5.368, p < 0.001, □^2^ = 0.392) suggesting different 3D-MOT performance across time between groups. There was no interaction on accuracy or RT. To clearly identify these effects, post-hoc comparisons are outlined in the sub-sections to follow.

The iMOT, Combi and Consol groups significantly improved their speed thresholds in ST_MOT_ and DT conditions between sessions T01 and T12 (all p < 0.05) and between T12 > T06 > T01 in both conditions for the Consol group only (see **supplementary material** for statistical values). Importantly, the slope analysis revealed that learning rates were significantly superior in the Consol compared to the Combi group during the DT task (U = 4.00, p = 0.003; Figure 5). In the iDM group, there was a significant difference between T12 > T01 in the ST_MOT_ condition which could most probably be attributed to a practice effect on 3D-MOT due to task exposure during the three evaluation sessions (T01, T06, T12). In fact, the first sessions of exposure to 3D-MOT are characteristic of a greater improvement (*e.g.* Faubert 2013). The iDM group significantly improved in accuracy during ST_DM_ between sessions T01 and T12 (p = 0.025; 25 % improvement). There was also a significant improvement of about 8 % in accuracy in the Consol group between sessions T06 and T12 (p = 0.026) in ST_DM_. Similarly, the improvement in the Combi group was also about 8 % but between sessions T01 and T12 (p = 0.051). There was no significant difference in RT between sessions and in any of the groups and conditions.

The one-way ANOVAs revealed a significant difference on speed thresholds between *Groups* at session T12 in the ST_MOT_ (F[3, 28] = 3.600, p = 0.027, □^2^ = 0.302) and DT (F[3, 28] = 10.667, p < 0.001, □^2^ = 0.561) conditions. Multiple comparisons demonstrated that only the Consol group was superior to the iDM group in STMOT (p = 0.021) and DT conditions (p < 0.001). Combi (p = 0.002) and iMOT (p = 0.035) groups were superior to iDM in DT condition. There was no significant difference between *Groups* on accuracy and RT.

**Figure 5.**
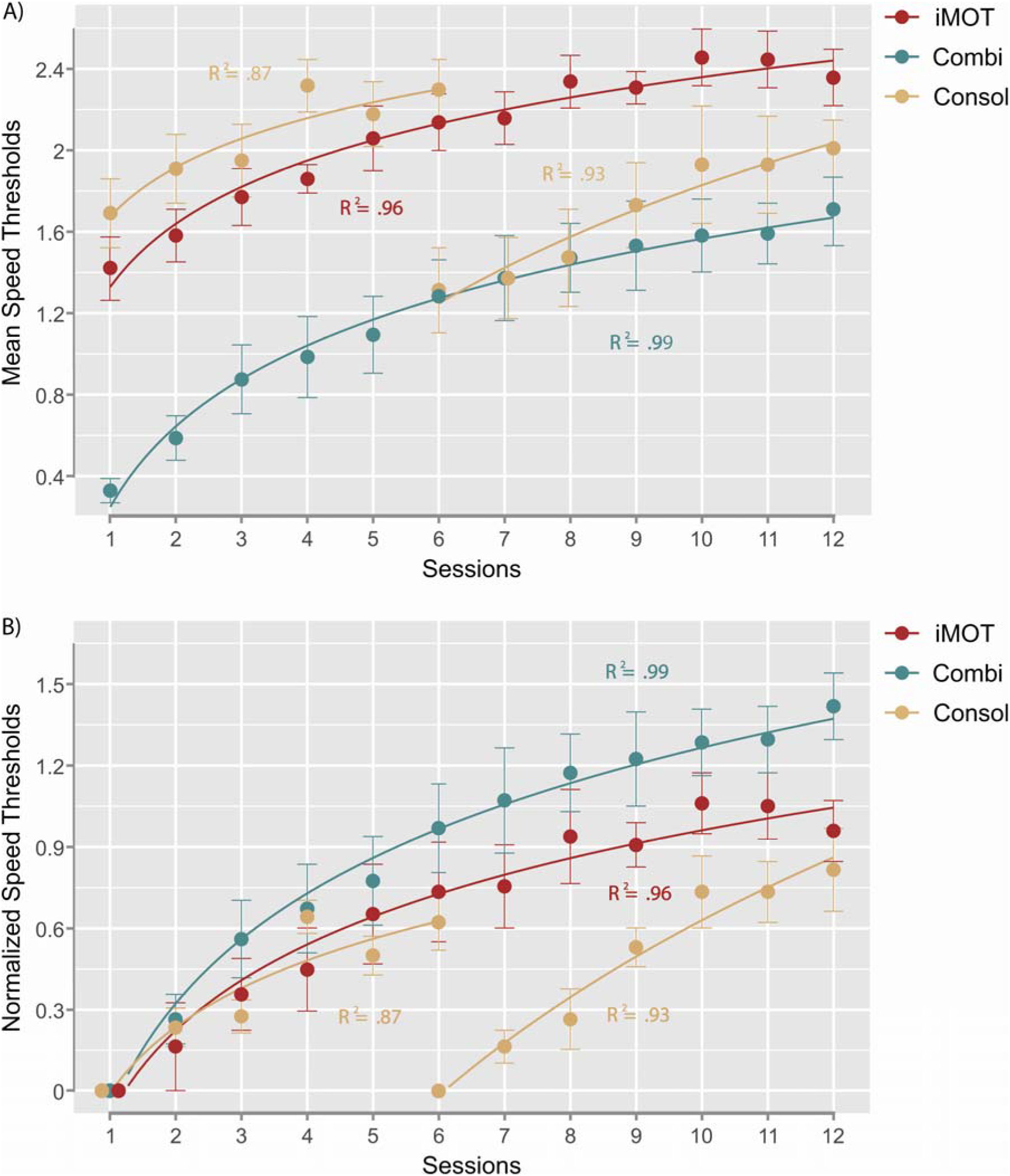
3D-MOT curves of A) mean speed thresholds and B) normalized speed thresholds across sessions. The fits shown are logarithmic regression functions and the R^2^ corresponds to the amount of variance explained by the fit. Error bars represent SEM.

## Discussion

Firstly, the results of this experiment indicated an important dual-task cost caused by the combination of 3D-MOT with a motor decision-making task. The additional task generated an important decrease in 3D-MOT speed thresholds. The results also demonstrated, however, that training reduced the difference between dual-task conditions and single-task conditions on 3D-MOT performance such that the dual-task condition began to converge toward the single-task condition, in particular in participants who followed the consolidation training regimen (Consol group). There was no apparent impact of dual-tasking on the motor task accuracy, but rather on RT, reflecting that the workload generated by the combined paradigm did not alter the quality of the execution and instead affected time execution.

Another important finding regarding the training methodology was that 3D-MOT performance and birdie interception accuracy were improved following training. More specifically, the consolidation training regimen seemed to give the best improvement across time on 3D-MOT performance and also revealed faster improvement on the motor decision-making task compared to the combined training. Importantly, the 3D-MOT learning rate in a dual-task condition was superior following a training regimen including a consolidation phase, rather than following simultaneous tasks’ combination. According to the present results, consolidated training should be prioritized when combining 3D-MOT with a motor task as previously experimented by Quevedo and colleagues (Quevedo, Blázquez et al. 2015).

In terms of the potential implication for perceptual-cognitive training, the findings revealed that the combined 3D-MOT methodology produced high challenge training conditions that are also highly trainable. In addition, the results suggested that consolidation training should be favoured when combining 3D-MOT with an additional motor execution task. The combined 3D-MOT motor exercise represents an interesting form of training by including a complex cognitive component to simple or more specific physical exercises. It acts on the coupling between perception, cognition and action which better reflects the reality of the sport environment. It would also be interesting to explore if execution accuracy and processing speed can be better trained with other execution movements such as when there is a more contextual judgement to make (*i.e.* attacking) than only anticipating the reception of the birdie (*i.e.* defensive action). The paradigm could be a suitable training program before the return to play in early season or after an injury, when athletes begin to engage physically. Further research will be needed to assess the transfer capability of this combined paradigm on sport performance.

## Experiment 2

The dynamic environment surrounding us requires to integrate an important number of visual information in order to make appropriate decisions. For example, navigating into a crowd requires to spread our attention on the dynamic information to orientate our actions and reach our goals; at the same time, it also requires us to focus on movements from others to avoid collisions. In sports, it is necessary to be able to perceive a number of elements in the dynamic scene and at the same time to pick up relevant information (*e.g.* body langage from a direct opponent) to execute the most appropriate decision or action. Thus, the second experiment intended to evaluate a dual-task paradigm built to mimic the important perceptual-cognitive load imposed during the previous situations. A life-sized combination of 3D-MOT with a embedded perceptual decision-making task was used. The biological motion perception (BMP) technique was employed for the perceptual exercise. Biological motion perception represents the capacity to recognize the kinematic presentation of movements reduced to a few moving dots placed on the major joints of the body (Johansson 1973). This representation allows human observers to recognize complex actions spontaneously from various animations such as walking. The paradigm allows to determine what the observer perceives solely on kinematics, while other motion cues are eliminated (Sparrow and Sherman 2001). This task has been used to highlight athlete expertise in sport science, has shown potential for the study of spatial characteristics of perception related to sport action (Abernethy and Parker 1989, Ward, Williams et al. 2002, Wright, Bishop et al. 2011) and has allowed researchers to assess perception of sports action within a life-sized virtual environment using stereoscopic displays (Bideau, Kulpa et al. 2010, Romeas and Faubert 2015). In the present study, a virtual biological motion pattern was employed to assess users’ decision-making ability based on specific kinematics while also spreading their attention on the dynamic scene to visually track moving targets (3D-MOT task). Previous evidence has shown that peforming two visual-spatial attention tasks (*i.e* visual tracking and visual search) that share common resources is challenging, but not impossible, for the human brain (Alvarez, Horowitz et al. 2005). In addition, it was found that interference is greater when a visual search task (*e.g.* MOT) is combined with an additional visual task compared to when combined with a task in a different sensory modality (*e.g.* auditory, tactile) (Arrighi, Lunardi et al. 2011, Wahn and Konig 2016). This could suggest a difference in the outcome of Experiment 2 (perceptual task) compared to Experiment 1 (motor task).

Similarly to Experiment 1, we expect an important dual-task cost generated by the addition of the perceptual decision-making task. Based on previous evidence (Alvarez, Horowitz et al. 2005, Arrighi, Lunardi et al. 2011) and Experiment 1 findings, we anticipate a reasonable degree of performance on both tasks in addition to improvement that could be altered by the perceptual nature of the BMP task. Like Experiment 1, the second objective was to assess the impact of different training regimens on task improvement. To test these hypotheses, the same four training regimens as in Experiment 1 were used. All groups were evaluated on each task (single-task 3D-MOT, dual-task 3D-MOT, single decision-making task) at sessions 1, 6 and 12.

## Material and methods

### Participants

Twenty-six healthy young but non-athlete adults (8 women; 22.96 ± 2.9 years old; 5.46 ± 2.82 hours of weekly physical training of any sport) were recruited for the study. The same screening survey precautions and ethical considerations as in Experiment 1 were met.

### Apparatus

The same immersive virtual environment was used as in Experiment 1.

### Experimental setup

The same 3D-MOT task was used as in Experiment 1 (Figure 1). The BMP task (Figure 6), consisted of the discrimination of the walking direction (right or left) of a point-light walker (Romeas and Faubert 2015). The point-light walker (Legault, Allard et al. 2013, Romeas and Faubert 2015) was a dynamic representation of human forms and was made up of 15 black dots, which represented the head, upper and lower trunk, shoulders, hips, elbows, wrists, knees, and ankles on a white background. Each dot had a diameter of 0.1 m. The height of the point-light walker was 1.80 m displayed at a virtual distance of 4 m from the observer. The presentation lasted for 1 s and contained 30 frames. The inter-stimulus interval was between 500 and 1000 ms in a random design. Point-light walkers were presented walking leftward or rightward (*i.e.* forced choice paradigm). A constant stimuli procedure with random angles of presentation across trials was used (−15, −4, −2, 0, 2, 4 and 15° from front-facing). All of the angles were randomly presented twenty times in each of the three experimental blocks and their order of presentation varied according to the constant stimuli procedure. Participants performed the task while seated and their performance was assessed through their decision accuracy (in %) and their response speed (difference in seconds between the appearance of the point-light walker and the participants’ decision; *i.e.* reaction time; RT). These two scores corresponded to the mean of the scores obtained with each of the three angles (2, 4 and 15°).

**Figure 6.**
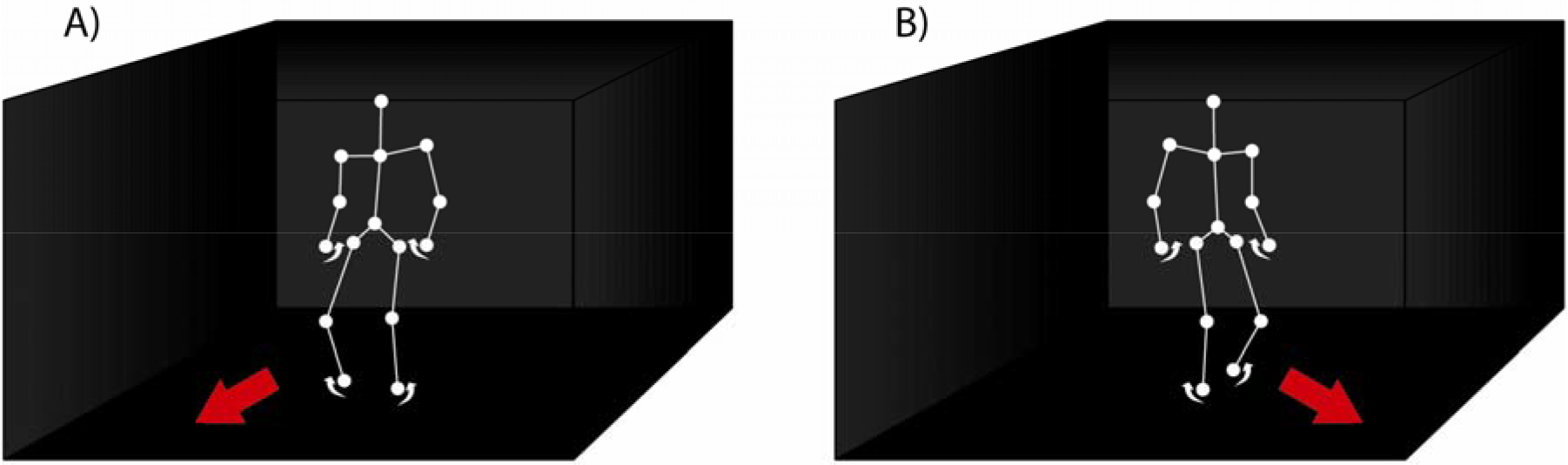
Illustration of the Biological Motion Perception task. The task consists of choosing whether an animated point-light walker is walking A) leftward or B) rightward from the subjects own vertical reference. In short, the task is to determine whether the walker’s predicted path will end up to the left or to the right of the subject’s own vertical center of reference. Red and white arrows as well as connecting white lines between dots are used here as a visual aid and were not presented during the experiment.

### Training regimen

The same training regimens were used as in Experiment 1 (Figure 3). *i)* Seven participants were engaged in the isolated 3D-MOT regimen (iMOT). *ii)* Seven participants were engaged in a dual-task training called combined (Combi). The combined task was a perceptual decision-making task (BMP). During each 8 s 3D-MOT trial, participants had to simultaneously perform 3 decision-making trials. *iii)* Six participants were involved in a training regimen that we called consolidation (Consol). *iv)* Finally, six participants performed the isolated decision-making task training (iDM).

### Protocol

The same protocol was used as in Experiment 1.

### Statistical analyses

The same procedure was used as in Experiment 1.

## Results

### Dual-task cost

There was a significant main effect of *Task condition* on speed thresholds (F[1, 22] = 289.635, p < 0.001, □^2^ = 0.929), on accuracy (F[1, 22] = 52.023, p < 0.001, □^2^ = 0.703) and on RT (F[1, 22] = 9.223, p = 0.006, □^2^ = 0.295) which demonstrated an important change in performance on all variables between the single and combined task.

### Training effectiveness across Sessions and between Groups

There was a main effect of *Sessions* on speed thresholds in ST_MOT_ (F[2, 44] = 51.138, p < 0.001, □^2^ = 0.699) and DT (F[2, 44] = 43.603, p < 0.001, □^2^ = 0.665), on accuracy in DT (F[2, 44] = 11.497, p < 0.001, □^2^ = 0.343) and on RT in STdm (F[2, 44] = 25.384, p < 0.001, □^2^ = 0.536) and DT (F[2, 44] = 4.887, p = 0.012, □^2^ = 0.182). While speed thresholds and RT performance were improved throughout sessions (all p < 0.017), pairwise comparisons performed on accuracy in DT demonstrated a decrease in accuracy between sessions T01 > T12 (p = 0.002) which was attributed to a speed-accuracy tradeoff (Figure 7). In fact, there was a speed-accuracy tradeoff on the perceptual decision-making task along the training in ST_DM_ (r_T01_ = 0.571, p = 0.002, r_T06_ = 0.543, p = 0.004, r_T12_ = 0.659, p < 0.001) and DT (r_T06_ = 0.468, p = 0.016, r_T12_ = 0.573, p = 0.002). There was no main effect of *Groups.* However, the ANOVA revealed a significant interaction between *Groups* and *Sessions* factors on speed thresholds in ST_MOT_ (F[6, 44] = 4.547, p = 0.001, □^2^ = 0.383) that was almost significant in DT (F[6, 44] = 2.092, p = 0.073, □^2^ = 0.222) which suggested differences in 3D-MOT performance across time between groups. The analysis also demonstrated a trend between *Groups* and *Sessions* factors on RT in DT (F[6, 44] = 2.182, p = 0.063, □^2^ = 0.229). To clearly identify these effects, post-hoc comparisons are outlined in the following sub-sections.

**Figure 7.**
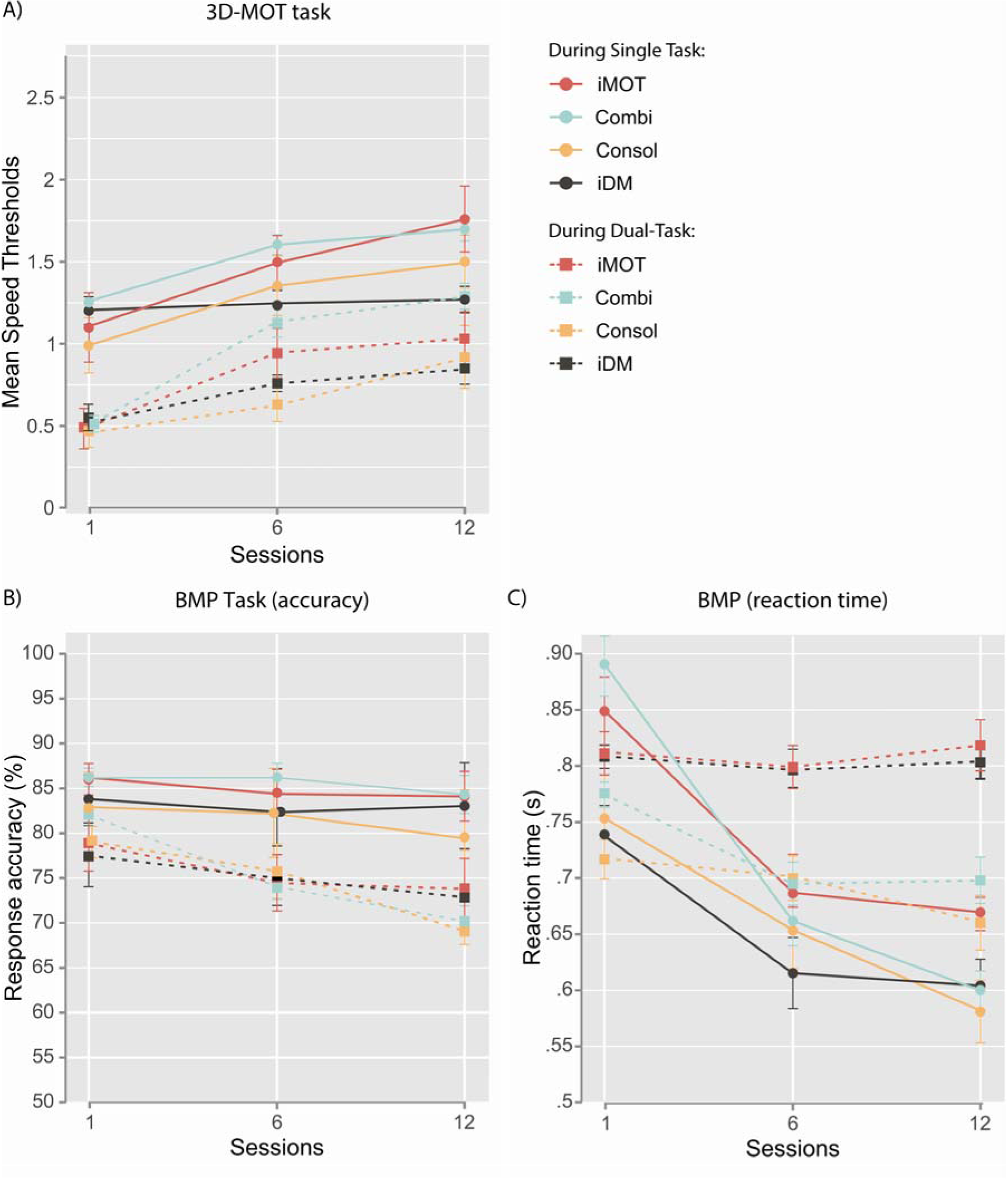
Training effectiveness across groups and sessions on the A) 3D-MOT and B), C) Decision-Making tasks (accuracy and reaction times, respectively). Circles and solid lines represent the performances obtained in single condition whereas squares and dashed lines correspond to performances recorded during the dual-task condition. Error bars represent SEM.

Similarly to Experiment 1, iMOT, Combi and Consol groups significantly improved their speed thresholds in ST_MOT_ and DT conditions between sessions T01 and T12 (all p < 0.05). However, in constrast to what was observed in Experiment 1, learning rates were not different between Combi and Consol groups (U = 18.000, p = 0.731; Figure 8). There was no substantial changes on speed thresholds in the iDM group. In addition, there was a significant decrease in accuracy in the Combi group in DT between T01 > T12 (all p < 0.05) which was caused by a prioritization of response speed over response accuracy (speed-accuracy tradeoff). In fact, RT diminished significantly in this condition between T01 and T12 in Combi (t[6] = 2.900, p = 0.027, □^2^ = 0.584). There was no meaningful effect of training on accuracy in STDM (all p > 0.05) in any group which suggest ceiling effect in the participtants’ capacity to accurately perceive the biological motion pattern. However, RT significantly decreased in all groups between T01 and T12 in ST_DM_ (all p < 0.05) which can be interpreted as a form of task improvement.

The one-way ANOVAs revealed only one significant difference between *Groups* on RT at session T12 in DT (F[3, 25] = 3.167, p = 0.045, □^2^ = 0.302) but multiple comparisons did not demonstrate any meaningful difference.

**Figure 8.**
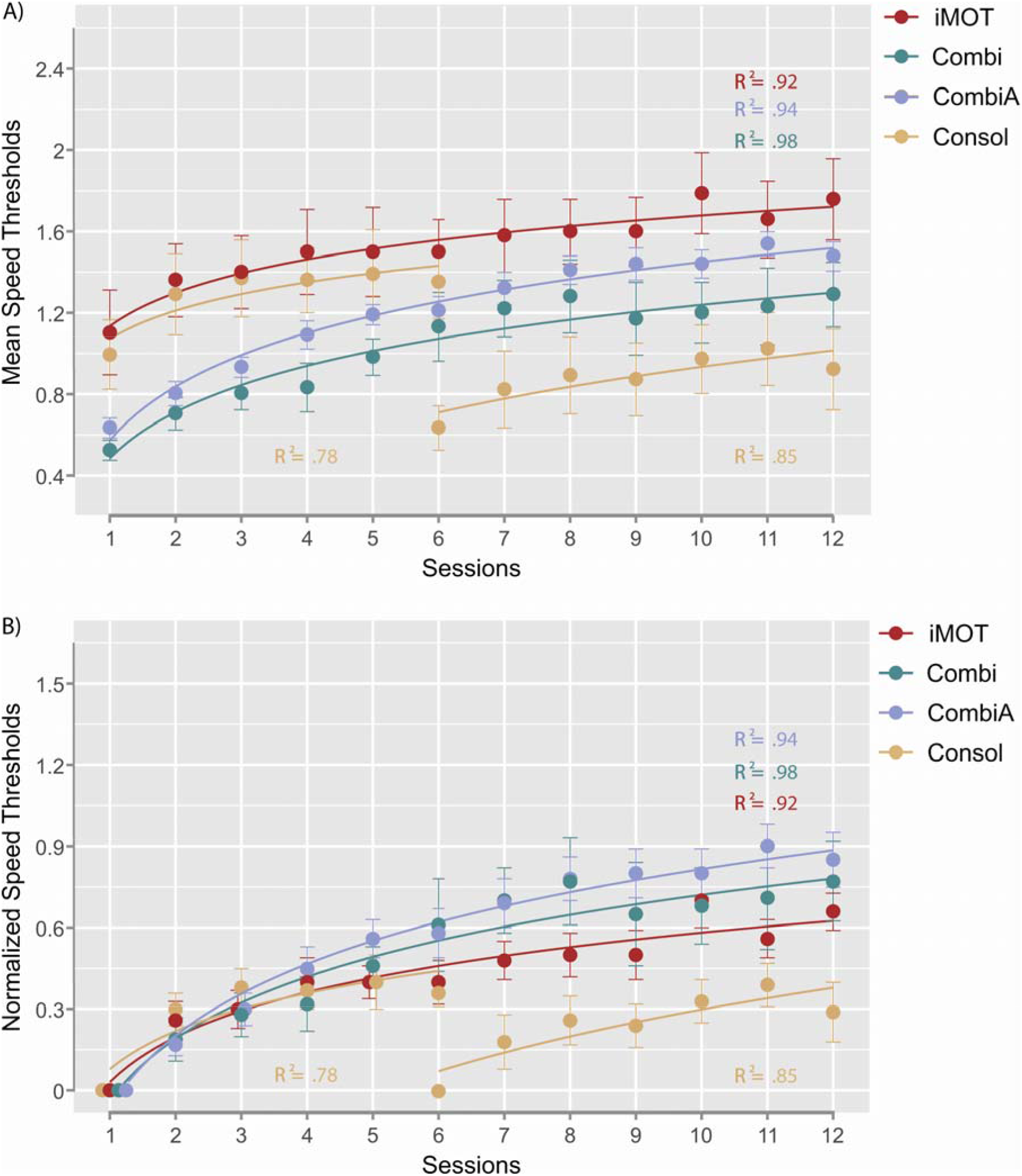
3D-MOT curves of A) mean speed thresholds and B) normalized speed thresholds across sessions. Combi refers to the results observed in non-atheltes participants whereas CombiA refers to the results obtained in athletes. The fits shown are logarithmic regression functions and the R^2^ corresponds to the amount of variance explained by the fit. Error bars represent SEM.

## Discussion

Like in Experiment 1, an important dual-task cost was observed when combining 3D-MOT with a decision-making task. However, the additional perceptual task was even more impacted during this experiment as evidenced by a dual-task cost on both the accuracy and RT of the BMP task. Contrary to Experiment 1, the training did not reduce the differences between dual-task conditions and single-task conditions on 3D-MOT performance. The results raise interesting questions regarding the allocation of attentional resources when combining two tasks from the same sensory modality (*e.g.* visual).

Despite highly challenging conditions (dual-task cost), task performance was executed beyond chance level and 3D-MOT as well as RT performance were generally improved following training. These findings supports previous evidence by Alvarez and colleagues (2005) showing that performing two visual-spatial tasks sharing common resources is challenging but not out of reach (Alvarez, Horowitz et al. 2005). Importantly, the perceptual decision-making task was prone to a speed-accuracy tradeoff (*i.e.* response speed was improved while response accuracy declined) which could be interpreted as a strategy to allocate more resources to the 3D-MOT task by performing the additional task more quickly. Moreover, contrary to Experiment 1, there was no evidence that dual-task training (combined or consolidated) offered higher dual-task performance,. Even the learning curves on 3D-MOT performance in dual-task conditions did not differ between the simultaneous (Combi) and consolidated (Consol) training regimens. The results observed on dual-task cost and on training accuracy tend to support other findings showing that performing two tasks with common sensory modalities (*e.g.* visual) leads to an important interference on task performance (Arrighi, Lunardi et al. 2011, Wahn and Konig 2016) and, now, considering the present results, on training effectiveness.

The nature of the additional task, which was purely perceptual in Experiment 2, might have led to the present results. But the difference observed between Experiment 1 and 2 could also be attributed to the nature of the participants (athletes vs non-athletes). Indeed, as athletes are more inclined to process multiple tasks at the same time (*e.g.* Chaddock, Neider et al. 2011), they might be better than mere novices to execute the dual-task paradigm as observed in Experiment 1. To test this hypothesis, we assessed whether the difference observed between Experiments 1 and 2 was due to the nature of the additional task (motor vs perceptual) or to presumed multiple tasks processing abilities. In this regard, we performed a repeated measures ANOVA (*Groups* x *Sessions*) on the six dependent variables using two groups. We compared Experiment 2 Combi group to sixteen university-level athletes (mean = 20.88 ± 2.03 (SD) years old; 12.56 ± 3.95 years of sport practice (soccer association and football); 12.75 ± 6.46 hours of workout per week) who subsequently participated in the simultaneous training regimen (Combi group) following exactly the same protocol and training conditions. Overall, we found no significant difference on any variable (speed thresholds, accuracy, RT; Figure 8 and **supplementary material**) between these two groups which supports the idea that using combined tasks sharing common sensory modalities interferes strongly with task performance and training, without regards to the athletic status of the participants.

Despite the interesting results raised by the present experiment, it is worth noting that the methodology used involved some limitations. In fact, combining 3D-MOT and BMP was influenced by a speed-accuracy tradeoff. Habituation to point-light pattern of movements produced faster responses with participants becoming more confident with their choices. And because no real-time feedback was given regarding the decision made, the chances to make repetitive mistakes were higher. As interpreted earlier, the speed-accuracy tradeoff could also result from a strategy to allocate more resources to perform the 3D-MOT task in dual-task conditions. In the future, it would be necessary to add feedback to allow the user to recognize when he is right or wrong. Another solution would be to diversify the nature of the point-light patterns of movement (*i.e.* multiple sports-related movements) to favor participants’ engagement to the task and potentially avoid ceiling effects on accuracy. These advancements would eventually improve the task methodology and consequently raise interest for this perceptual-cognitive training. In turn, the paradigm could have very practical uses in cases where athletes are unable to physically respond, acting as a form of observational learning and cognitive stimulation.

## General discussion

The present study assessed the conception of a multitasking perceptual-cognitive training exercise including 3D-MOT combined with decision-making tasks from different nature. Four training regimens were tested during single and dual-task conditions throughout 3 evaluation sessions (T01, T06 and T12). Trained groups included the isolated 3D-MOT task (iMOT), 3D-MOT simultaneously combined with a decision-making task (Combi), 3D-MOT combined with a decision-making task following a 3D-MOT consolidation phase (Consol) and the isolated decision-making task (iDM). Importantly, the additional decision-making task was either motor (Experiment 1, birdie interception) or perceptual (Experiment 2, BMP).

The main findings of this study indicate that the combined perceptual-cognitive training paradigm represents a high-level challenging environment, as illustrated by an important interference (dual-task cost) on task performance under dual-task conditions. Importantly, despite the difficulty inherent to the dual-task exercise, task performance was trainable which supports the potential implication of this technique for perceptual-cognitive training. Secondly, the four different training regimens (iMOT, Combi, Consol, iDM) did not equally influence the performance exhibited in each condition (ST_MOT_, ST_DM_, DT) with dual-task trained groups tending to show the best improvements. This was especially true for speed threshold scores and particularly the consolidation training group (Consol) when 3D-MOT was combined with a motor task (Experiment 1). However, when 3D-MOT was combined with a perceptual task sharing common attentional resources, consolidation and combined training groups did not exhibit different levels of performance (Experiment 2).

The first objective of the study was to validate whether combining 3D-MOT with a decision-making exercise could offer more challenging conditions while integrating broader contextual information. The dual-task condition produced a significant reduction in task performance in Experiment 1 and 2 which is consistent with most of the previous studies using dual-task MOT, 3D-MOT or spatial visual tests (Alvarez, Horowitz et al. 2005, Allen, McGeorge et al. 2006, Zhang, Xuan et al. 2010, Thomas and Seiffert 2011, Faubert and Sidebottom 2012, Quevedo, Blázquez et al. 2015, Lapierre, Cropper et al. 2017). For example, Thomas & Seiffert (2011) observed a decrease in MOT performance between single and dual-task conditions (while walking) under high tracking load (≥3 targets). Likewise, Zhang and colleagues have concluded clear evidence that when a MOT task and a visual working memory task are performed concurrently, both tasks are disrupted as long as either task is sufficiently challenging to consume the resources available (Zhang, Xuan et al. 2010, Lapierre, Cropper et al. 2017). Interestingly, comparing Experiment 2 to Experiment 1 highlighted a greater dual-task cost when the additional task shared similar sensory modalities with the primary task. Previously, Arrighi and colleagues (2011) found that a MOT task selectively interfered with a visual discrimination task sharing common attentional resources, compared to an auditory discrimination task (Arrighi, Lunardi et al. 2011). In their review, Wahn and Konig (2017) also supported that simultaneously performing a visual spatial attention task and an object-based attention task recruits partially shared attentional resources which can explain a superior interference compared to when tasks are from different sensory modalities (Wahn and Konig 2017). Based on this evidence, it is suggested that the perceptual nature of the combined task in Experiment 2 was responsible for the greater interference compared to Experiment 1 (motor decision-making task). To our knowledge, the present study is the first comparison of the simultaneous processing of both MOT and BMP tasks. The BMP task is known to induce a selective activation of the brain, especially in the superior temporal sulcus, a region that is part of the action observation network (Oram and Perrett 1994, Vaina, Solomon et al. 2001, Ptito, Faubert et al. 2003). On the other hand, the MOT task has been found to activate a certain number of higher brain areas involved in attentional processes such as parietal and frontal regions of the cortex as well as the middle temporal complex (Culham, Brandt et al. 1998, Howe, Horowitz et al. 2009). Shared resources between these systems were potentially solicited during the execution of the BMP and 3D-MOT tasks but further research, beyond the scope of the present study, will be needed to firmly conclude this hypothesis.

Another interesting finding showed that although the virtual stimulation provided throughout the dual-task paradigm was costly in terms of central attention, performance and learning were not totally hampered in Experiment 1 and 2. Indeed, participant’s speed thresholds, accuracy and RT were still producing acceptable levels of efficiency under dual-task conditions, whether it was motor or perceptual, reflecting that the exercices were processed beyond level of chance (mean accuracy thresholds always just above a noticeable difference [> 50%] and response time to decision-making in accordance with previous evidence; *e.g.* Land and McLeod 2000, Romeas and Faubert 2015). For example, there is previous evidence where 3D-MOT combined with high cognitive load conditions (*i.e.* flying a jet) could not be performed with such accuracy (Hoke, Reuter et al. 2017). In addition, Alvarez and colleagues (2005) demonstrated that while the decrease in performance during dual-tasking was attributed to a central executive cost, two visual-spatial attentional tasks were not continuously relying on the same cognitive resources (Alvarez, Horowitz et al. 2005). Interestingly, the authors combined visual tracking (MOT) and visual search tasks. Their study suggested that it is possible to search through and track spatially overlapping sets of stimuli although there were significant dual-task costs which is congruent with Experiment 2. The present study confirms that it is possible to visually track and search through overlapping virtual elements in addition to taking a real-time decision in response to a specific cue.

Moreover, the results demonstrated that performance in the life-sized virtual perceptual-cognitive paradigm is highly trainable. First of all, Experiment 1 tended to support the benefits of dual-task training for task improvement. Indeed, dual-task training has shown some evidence of transfer benefits over single-task training in more general domains (Silsupadol, Shumway-Cook et al. 2009, Theill, Schumacher et al. 2013, Desjardins-Crepeau, Berryman et al. 2016). Importantly, Laufer (2008) demonstrated the importance of using challenging cognitive tasks in dual-task training: their participants obtained better postural control when the balance training was coupled with a highly demanding cognitive task (arithmetic manipulation) compared to a cognitive task that required little attention (e.g. forward counting; Laufer 2008). Moreover, skill retention was enhanced in the group trained under conditions with greater attentional demands. Gabbett and colleagues (2011) have investigated the effects of single-task and dual-task training on the acquisition, retention, and transfer of skill (drawing and passing) in high performance rugby league players (Gabbett, Wake et al. 2011). Dual-task training did not significantly outperform single-task training, but improved the dual-task performance of the draw and pass skill. In Experiment 1, although 3D-MOT task performance indicated a reliable difference between dual-task and single-task after training, there was a decrease in the magnitude of the difference after training when 3D-MOT was combined with a motor task such that the dual-task condition began to converge toward the single-task condition. It is suggested that the cortical processors involved in dual-task performance became more adept at a process more specific to the dual-task condition, such as processing 3D-MOT while making a decision (Erickson, Colcombe et al. 2007). Studies have also shown a practice effect on dual-task performance, suggesting that dual-task performance relies on attentional control strategies. This implies that training and learning an optimal strategy could help improve dual-task performance (Bherer, Kramer et al. 2008). Other evidence has demonstrated that reduction in dual-task costs require practice on the combination of the two updating tasks, not just practice on an individual task (Oberauer and Kliegl 2004). A fundamental objective of perceptual-cognitive training is the reduction of the attentional resources required to produce a particular outcome. It is possible that participants had learnt an optimal strategy throughout the motor combined 3D-MOT training to a point where the dual-task cost was decreased compared to participants trained with single tasks.

On the other hand, results observed in Experiment 2 suggested that dual-task training do not offer a meaningful advantage when both tasks are from the same sensory modality. As described previously, performing two tasks with common sensory modalities (*e.g.* visual) leads to an important interference on task performance (Arrighi, Lunardi et al. 2011, Wahn and Konig 2017). The results observed in Experiment 2 seem to support that this type of combination impacts training effectiveness as well, compared to when the two tasks are from different sensory modalities (Experiment 1).

The present findings also provide important insights regarding training regimens’ efficiency within the context of dual-task training. When combining 3D-MOT with a motor decision-making task (Experiment 1), the consolidation training regimen gave the best improvement across time on 3D-MOT performance and showed similar benefits as the simultaneous training group (Combi) with less dual-task training time. More importantly, 3D-MOT learning rate in dual-task conditions was superior in the Consol group which first learnt how to process 3D-MOT in isolation (phase 1) before engaging in the dual-task training (phase 2). These results raised the importance of consolidated training before the engagement in simultaneous training for rapid performance enhancement, especially if a variety of skills have to be mastered (Quevedo, Blázquez et al. 2015). A few studies have tried to compare sequential and simultaneous trainings; for a methodological example, see the ongoing work by Lee and colleagues (2016) in patients with mild cognitive impairment (Lee, Wu et al. 2016). In the sport domain, there is very few comparisons between the simultaneous and sequential approaches. Recently, video-based training has been used to demonstrate that presenting tennis actions or shots in the order they occur in the performance environment led to better performance in the field-based transfer test compare to a non-sequential trained group (Broadbent, Ford et al. 2017). The authors explained that sequential structure of practice potentially improves contextual information. The results of the present study suggest that consolidated training might be more beneficial than simultaneous training when combining perceptual-cognitive and motor exercises (*i.e.* birdie interception). In other words, initial familiarization with the unpredictable dynamic environment is suggested to be an important requirement for higher upcoming motor dual-tasking performance.

Finally, it has been recently proposed that multiple sources of information other than simple visual kinematic cues picked-up from the direct opponent, play a significant role in fast judgements (de Oliveira, Lobinger et al. 2014, Canal-Bruland and Mann 2015). These sources of contextual factors are present inside the unpredictable dynamic sport scenario and must be taken into account for better and faster decision making. The training tool used in the present study seeks to approach broader ecological and contextual situations by embedding a perception-action component (decision-making task) in a purely perceptual-cognitive task (3D-MOT) that requires to process complex and unpredictable dynamic visual scenes. For the first time, this paradigm proposes to integrate components supported by both the sport specific and the cognitive component approaches (Alves, Voss et al. 2013) during a perceptual-cognitive training. Moreover, by testing different kinds of training conditions, this study provides important insights on the requirements for building training regimens with a better ratio between time investment and efficiency. In the future, the transfer effectiveness of this new perceptual-cognitive training methodology needs to be put to the test to assess whether broader contextual paradigms are producing superior outcomes for sports performance.

## Acknowledgments

The authors would like to thank Stéphanie Ferland and Philippe Letendre-Joachim for their great work with testing participants. We would also like to thank Satya Ortiz-Gagné and Vadim Sutyushev for programming the tasks. We thank Robyn Lahiji and Jesse Michaels for their help in editing the manuscript. Special thanks to the participants.

## References

Abernethy B. and S. Parker (1989). Perceiving joint kinematics and segment interactions as a basis for skilled anticipation in squash. Proceedings of the 7th World Congress in Sport Psychology. C. K. Giam, K. K. Chook and K. C. The. Singapore, International Society of Sport Psychology: 56–58.

Allen, R., P. McGeorge, D. G. Pearson and A. Milne (2006). “Multiple-target tracking: A role for working memory?” Quarterly journal of experimental psychology 59(6): 1101–1116.

Alvarez, G. A., T. S. Horowitz, H. C. Arsenio, J. S. DiMase and J. M. Wolfe (2005). “Do Multielement Visual Tracking and Visual Search Draw Continuously on the Same Visual Attention Resources?” Journal of Experimental Psychology: Human Perception and Performance 31(4): 643–667.

Alves, H., M. W. Voss, W. R. Boot, A. Deslandes, V. Cossich, J. I. Salles and A. F. Kramer (2013). “Perceptual-cognitive expertise in elite volleyball players.” Frontiers in Psychology 4: 36.

Arrighi, R., R. Lunardi and D. Burr (2011). “Vision and Audition Do Not Share Attentional Resources in Sustained Tasks.” Frontiers in Psychology 2.

Bherer, L., A. F. Kramer, M. S. Peterson, S. Colcombe, K. Erickson and E. Becic (2008). “Transfer effects in task-set cost and dual-task cost after dual-task training in older and younger adults: further evidence for cognitive plasticity in attentional control in late adulthood.” Experimental aging research 34(3): 188–219.

Bideau, B., R. Kulpa, N. Vignais, S. Brault, F. Multon and C. Craig (2010). “Using virtual reality to analyze sports performance.” IEEE Computer Graphics and Applications 30(2): 14–21.

Broadbent, D. P., J. Causer, A. M. Williams and P. R. Ford (2015). “Perceptual-cognitive skill training and its transfer to expert performance in the field: future research directions.” European Journal of Sport Science 15(4): 322–331.

Broadbent, D. P., P. R. Ford, D. A. O’Hara, A. M. Williams and J. Causer (2017). “The effect of a sequential structure of practice for the training of perceptual-cognitive skills in tennis.” PLoS One 12(3): e0174311.

Canal-Bruland R. and D. L. Mann (2015). “Time to broaden the scope of research on anticipatory behavior: a case for the role of probabilistic information.” Frontiers in Psychology 6: 1518.

Chaddock, L., M. B. Neider, M. W. Voss, J. G. Gaspar and A. F. Kramer (2011). “Do athletes excel at everyday tasks?” Medicine & Science in Sports & Exercise 43(10): 1920–1926.

Culham, J. C., S. A. Brandt, P. Cavanagh, N. G. Kanwisher, A. M. Dale and R. B. Tootell (1998). “Cortical fMRI activation produced by attentive tracking of moving targets.” Journal of Neurophysiology 80(5): 2657–2670.

de Oliveira, R. F., B. H. Lobinger and M. Raab (2014). “An adaptive toolbox approach to the route to expertise in sport.” Frontiers in Psychology 5: 709.

Desjardins-Crepeau, L., N. Berryman, S. A. Fraser, T. T. Vu, M. J. Kergoat, K. Z. Li, L. Bosquet and L. Bherer (2016). “Effects of combined physical and cognitive training on fitness and neuropsychological outcomes in healthy older adults.” Clinical Interventions in Aging 11: 1287–1299.

Erickson, K. I., S. J. Colcombe, R. Wadhwa, L. Bherer, M. S. Peterson, P. E. Scalf, J. S. Kim, M. Alvarado and A. F. Kramer (2007). “Training-induced functional activation changes in dual-task processing: an FMRI study.” Cerebral Cortex 17(1): 192–204.

Faubert, J. (2013). “Professional athletes have extraordinary skills for rapidly learning complex and neutral dynamic visual scenes.” Scientific reports 3: 1154.

Faubert J. and L. Sidebottom (2012). “Perceptual-Cognitive Training of Athletes.” Journal of Clinical Sports Psychology(6): 85–102.

Gabbett, T., M. Wake and B. Abernethy (2011). “Use of dual-task methodology for skill assessment and development: examples from rugby league.” Journal of Sports Sciences 29(1): 7–18.

Harenberg, S., R. McCaffrey, M. Butz, D. Post, J. Howlett, K. D. Dorsch and K. Lyster (2016). “Can Multiple Object Tracking Predict Laparoscopic Surgical Skills?” Journal of Surgical Education.

Hoke, J., C. Reuter, T. Romeas, M. Montariol, T. Schnell and J. Faubert (2017). Perceptual-cognitive and physiological assessment of training effectiveness. Proceedings of the Interservice/ Industry Training, Simulation and Education Conference Orlando, FL.

Howe, P. D., T. S. Horowitz, I. A. Morocz, J. Wolfe and M. S. Livingstone (2009). “Using fMRI to distinguish components of the multiple object tracking task.” Journal of Vision 9(4): 10 11–11.

Johansson G. (1973). “Visual perception of biological motion and a model for its analysis.” Attention, Perception, & Psychophysics 14(2): 201–211.

Land M. F. and P. McLeod (2000). “From eye movements to actions: how batsmen hit the ball.” Nature Neuroscience 3(12): 1340–1345.

Lapierre, M. D., S. J. Cropper and P. D. L. Howe (2017). “Shared processing in multiple object tracking and visual working memory in the absence of response order and task order confounds.” PLoS One 12(4): e0175736.

Laufer Y. (2008). “Effect of cognitive demand during training on acquisition, retention and transfer of a postural skill.” Human Movement Science 27(1): 126–141.

Lee, Y. Y., C. Y. Wu, C. H. Teng, W. C. Hsu, K. C. Chang and P. Chen (2016). “Evolving methods to combine cognitive and physical training for individuals with mild cognitive impairment: study protocol for a randomized controlled study.” Trials 17(1): 526.

Legault, I., R. Allard and J. Faubert (2013). “Healthy older observers show equivalent perceptual-cognitive training benefits to young adults for multiple object tracking.” Frontiers in Psychology 4.

Legault I. and J. Faubert (2012). “Perceptual-cognitive training improves biological motion perception: evidence for transferability of training in healthy aging.” Neuroreport 23(8): 469–473.

Levitt H. (1971). “Transformed up-down methods in psychoacoustics.” The Journal of the Acoustical Society of America 49(2): Suppl 2:467+.

Mangine, G. T., J. R. Hoffman, A. J. Wells, A. M. Gonzalez, J. P. Rogowski, J. R. Townsend, A. R. Jajtner, K. S. Beyer, J. D. Bohner, G. J. Pruna, M. S. Fragala and J. R. Stout (2014). “Visual tracking speed is related to basketball-specific measures of performance in NBA players.” The Journal of Strength & Conditioning Research 28(9): 2406–2414.

Memmert D. (2009). “Pay attention! A review of visual attentional expertise in sport.” International Review of Sport and Exercise Psychology 2(2): 119–138.

Michaels, J., R. Chaumillon, D. Nguyen-Tri, D. Watanabe, P. Hirsch, F. Bellavance, G. Giraudet, D. Bernardin and J. Faubert (2017). “Driving simulator scenarios and measures to faithfully evaluate risky driving behavior: A comparative study of different driver age groups.” PLoS One 12(10): e0185909.

Oberauer K. and R. Kliegl (2004). “Simultaneous cognitive operations in working memory after dual-task practice.” Journal of Experimental Psychology: Human Perception and Performance 30(4): 689–707.

Oram M. W. and D. I. Perrett (1994). “Responses of Anterior Superior Temporal Polysensory (STPa) Neurons to “Biological Motion” Stimuli.” Journal of Cognitive Neuroscience 6(2): 99–116.

Parsons, B., T. Magill, A. Boucher, M. Zhang, K. Zogbo, S. Berube, O. Scheffer, M. Beauregard and J. Faubert (2016). “Enhancing Cognitive Function Using Perceptual-Cognitive Training.” Clinical EEG and Neuroscience 47(1): 37–47.

Pothier, K., N. Benguigui, R. Kulpa and C. Chavoix (2015). “Multiple Object Tracking While Walking: Similarities and Differences Between Young, Young-Old, and Old-Old Adults.” The journals of gerontology. Series B, Psychological sciences and social sciences 70(6): 840–849.

Ptito, M., J. Faubert, A. Gjedde and R. Kupers (2003). “Separate neural pathways for contour and biological-motion cues in motion-defined animal shapes.” NeuroImage 19(2 Pt 1): 246–252.

Quevedo, L., A. P. Blázquez, J. Solé i Fortó and G. C. Torradeflot (2015). “Perceptual-cognitive Training with the NeuroTracker 3D-MOT to Improve Performance in Three Different Sports.” Educació Física i Esports 119: 97–108.

Romeas T. and J. Faubert (2015). “Soccer athletes are superior to non-athletes at perceiving soccer-specific and non-sport specific human biological motion.” Frontiers in Psychology 6: 1343.

Romeas, T., A. Guldner and J. Faubert (2016). “3D-Multiple Object Tracking training task improves passing decision-making accuracy in soccer players.” Psychology of Sport and Exercise 22: 1–9.

Silsupadol, P., A. Shumway-Cook, V. Lugade, P. van Donkelaar, L. S. Chou, U. Mayr and M. H. Woollacott (2009). “Effects of single-task versus dual-task training on balance performance in older adults: a double-blind, randomized controlled trial.” Archives of Physical Medicine and Rehabilitation 90(3): 381–387.

Sparrow W. A. and C. Sherman (2001). “Visual expertise in the perception of action.” Exercise and Sport Sciences Reviews 29(3): 124–128.

Theill, N., V. Schumacher, R. Adelsberger, M. Martin and L. Jancke (2013). “Effects of simultaneously performed cognitive and physical training in older adults.” BMC Neuroscience 14: 103.

Thomas L. and A. Seiffert (2011). “How Many Objects are You Worth? Quantification of the Self-Motion Load on Multiple Object Tracking.” Frontiers in Psychology 2(245).

Vaina, L. M., J. Solomon, S. Chowdhury, P. Sinha and J. W. Belliveau (2001). “Functional neuroanatomy of biological motion perception in humans.” Proceedings of the National Academy of Sciences of the United States of America 98(20): 11656–11661.

Vartanian, O., L. Coady and K. Blackler (2016). “3D multiple object tracking boosts working memory span: Implications for cognitive training in military populations.” Military Psychology 28(5): 353–360.

Wahn B. and P. Konig (2016). “Attentional Resource Allocation in Visuotactile Processing Depends on the Task, But Optimal Visuotactile Integration Does Not Depend on Attentional Resources.” Frontiers in Integrative Neuroscience 10: 13.

Wahn B. and P. Konig (2017). “Is Attentional Resource Allocation Across Sensory Modalities Task-Dependent?” Advances in Cognitive Psychology 13(1): 83–96.

Ward, P., A. M. Williams and S. J. Bennett (2002). “Visual search and biological motion perception in tennis.” Research Quarterly for Exercise and Sport 73(1): 107–112.

Wright, M. J., D. T. Bishop, R. C. Jackson and B. Abernethy (2011). “Cortical fMRI activation to opponents’ body kinematics in sport-related anticipation: expert-novice differences with normal and point-light video.” Neuroscience Letters 500(3): 216–221.

Zhang, H., Y. M. Xuan, X. L. Fu and Z. W. Pylyshyn (2010). “Do objects in working memory compete with objects in perception?” Visual Cognition 18(4): 617–640.

